# Mapping the transcriptional regulatory network of a fungal pathogen by exploiting transcription factor perturbation

**DOI:** 10.1101/2025.05.10.653149

**Authors:** Dhoha Abid, Holly Brown, Chase Mateusiak, Tamara L. Doering, Michael R. Brent

## Abstract

*Cryptococcus neoformans* is a deadly fungal pathogen. Upon entering a mammalian host, it deploys a voluminous polysaccharide capsule that is necessary for it to survive host defenses and maintain an infection. Capsule expansion is regulated transcriptionally, as deletion of many transcription factors (TFs) alters capsule. Thus, we set out to map the transcriptional regulatory network of *C. neoformans* – that is, to identify the TFs that directly regulate each gene in the genome. First, we carried out RNA-seq of 120 single-TF-deletion strains, together with wild-type controls. We then applied NetProphet3, a TF network mapping algorithm, to predict the direct functional targets of each TF. Unexpectedly, analysis of this network indicated that there are no TFs that primarily regulate genes involved in capsule formation. Rather, the TFs that play a role in deploying capsule also regulate many other genes and processes. Comparison to a TF network map we built for *Saccharomyces cerevisiae*, a distantly related model yeast, identified pairs of TFs that are functionally orthologous – that is, their targets are enriched for orthologous genes. In many cases, these pairs are different from the ones identified by sequence homology alone. We suggest that network analyses should be used to complement sequence comparison when searching for functionally orthologous transcription factors. Our network map can be searched and visualized at http://cryptococcus.net.

## Introduction

*Cryptococcus neoformans* is a fungal pathogen that occurs in the environment and may infect mammals, including humans and mice. When infectious particles of *C. neoformans* are inhaled, they can initiate a lung infection. In immunocompetent individuals this is generally cleared, although latent infection may occur. In severely immunocompromised individuals, however, the infection persists in the lungs and frequently disseminates to the brain, where it causes potentially lethal meningitis (Meya and Williamson, 2024; Tugume, et al., 2023). *C. neoformans* is estimated to be the second most common cause of AIDS-related mortality (Rajasingham, et al., 2017) and responsible for 147,000 deaths annually (Denning, 2024; Rajasingham, et al., 2017).

*C. neoformans* is a basidiomycete, belonging to a phylum that diverged approximately 400 million years ago from the ascomycete lineage, which includes the model yeast *Saccharomyces cerevisiae* and other fungal pathogens like *Candida* and *Aspergillus* (Taylor and Berbee, 2006). It is distinguished by a large polysaccharide capsule that surrounds the cell and is central to its virulence (Boodwa-Ko and Doering, 2024). Upon entry into a host or exposure to conditions that mimic the host environment, the capsule expands considerably (Agustinho, et al., 2018; Wang, et al., 2018). Acapsular cells are avirulent (Fromtling, et al., 1982; Kwon-Chung and Rhodes, 1986), while larger capsules increase resistance to engulfment by phagocytes during infection (Bulmer and Sans, 1967; Kozel, 1977; Srikanta, et al., 2011).

We know that capsule production is regulated at the level of transcription (Gish, et al., 2016; Granger, et al., 1985), yet our understanding of this regulation is limited. One way to define regulatory processes is to generate a transcription factor (TF) network map (Bonneau, et al., 2006; Greenfield, et al., 2010; Huynh-Thu and Geurts, 2018; Jackson, et al., 2020; Madar, et al., 2010; Margolin, et al., 2006; Roy, et al., 2013; Siahpirani and Roy, 2017). Such maps describe the relationships between each TF in the genome and the genes that it regulates. A map consists of nodes and edges: nodes represent TFs and genes, and an edge exists between a TF and a target gene if the TF regulates that gene. Edges are described as: TF-target.

Many gene regulatory networks are intended to represent functional regulation rather than molecular interactions. In other words, edges indicate relationships between genes (or TFs and genes) that could be either direct or indirect. Here, we aim to generate a map that represents relationships in C*. neoformans* that are both direct and functional. In other words, the TF must physically bind to the regulatory DNA of the target gene (so it is direct), and as a result, it must modulate the transcription rate of that target (so it is functional). Importantly, this is not always the case: genes whose promoters are bound by a TF in ChIP-Seq or other binding location assays are frequently unaffected by deleting or over-expressing the TF, which shows that direct interaction does not always indicate a functional relationship (Kang, et al., 2022; Mahendrawada, et al., 2025).

To generate our network, we used NetProphet3, a state-of-the-art method for mapping TF networks that we recently developed (Abid and Brent, 2023). NetProphet3 uses as input gene expression profiles of cells with or without perturbation of a TF by, for example, deletion or overexpression. Because they are based on expression data, NetProphet3 maps are functional, in the sense that TFs are likely to affect the expression of their predicted target genes. Here, we show that the functional TF-target relationships predicted by NetProphet3 in *Cryptococcus* are highly enriched for direct interactions, consistent with our findings in *Saccharomyces cerevisiae* (Abid and Brent, 2023). Indeed, we show that NetProphet3 predicts the probability that a TF will bind a target gene in a ChIP-seq experiment with reasonable accuracy, using only gene expression data as input.

In this work, we generated a comprehensive dataset of gene expression changes in response to TF perturbation in *C. neoformans*. Specifically, we carried out RNA-Seq on 120 TF deletion strains (selected as described below) and used the resulting data as input for NetProphet3 to create a TF network map. The map assigns scores to all possible TF-target interactions that represent our confidence in the existence of a direct, functional relationship. After validating this map, we used it to identify functional orthologs of Cryptococcus TFs in *S. cerevisiae*. We also searched the network for TFs that primarily regulate genes involved in capsule formation, but found none, suggesting that capsule is not deployed independently but rather as part of an arsenal of cellular defense and stress response systems. Our new TF perturbation response dataset is available from GEO (GSE297962), the network map we constructed from it may be found at https://doi.org/10.5281/zenodo.17193243, and a visualization is available at https://cryptococcus.net.

## Results

### Overview of the transcriptome

We generated gene expression data for all of the single-TF deletion strains that we could either generate (Maier, et al., 2015) or obtain from the Madhani *C. neoformans* deletion collection (Boucher, et al., 2024). We grew these 120 strains, each lacking one TF (Supplementary File S1), along with wild-type (WT) controls, in standard laboratory conditions for rich medium growth (YPD, room air, 30°C). We then transferred them to capsule-inducing conditions for 90 minutes and measured gene expression levels by RNA-Seq. The capsule-inducing conditions we used, which are designed to mimic the host environment, include mammalian tissue culture medium (DMEM), 37 °C, and 5% CO_2_; these conditions are known to result in cells with large capsules (Maier, et al., 2015). We chose a 90-minute time point to reflect early changes in transcription and because it has yielded insights into cryptococcal gene regulation in our previous studies (Gish, et al., 2016; Maier, et al., 2015).

To enable rigorous analyses, we carried out RNA-Seq in a strictly controlled manner and with multiple biological replicates for each TF deletion strain (average = 4, Fig. 1A, File S1). We also included WT control cultures along with TF deletion strains in each experiment, totaling 122 WT samples. Under our conditions, we detected expression of 6746 genes, which included 163 of the 165 TFs in *C. neoformans*. Figure 1B shows the distribution of expression levels for all TFs in WT cells, while Supplementary Figure S1A provides the values for individual TFs. The most highly expressed TF genes were *RIM101* (O’Meara, et al., 2014) and *BZP1* (Jung, et al., 2015). Most TFs were measurably expressed in capsule-inducing conditions (Figure 1B), so we expected that deleting the corresponding genes could alter the expression of their targets.

**Fig 1.**
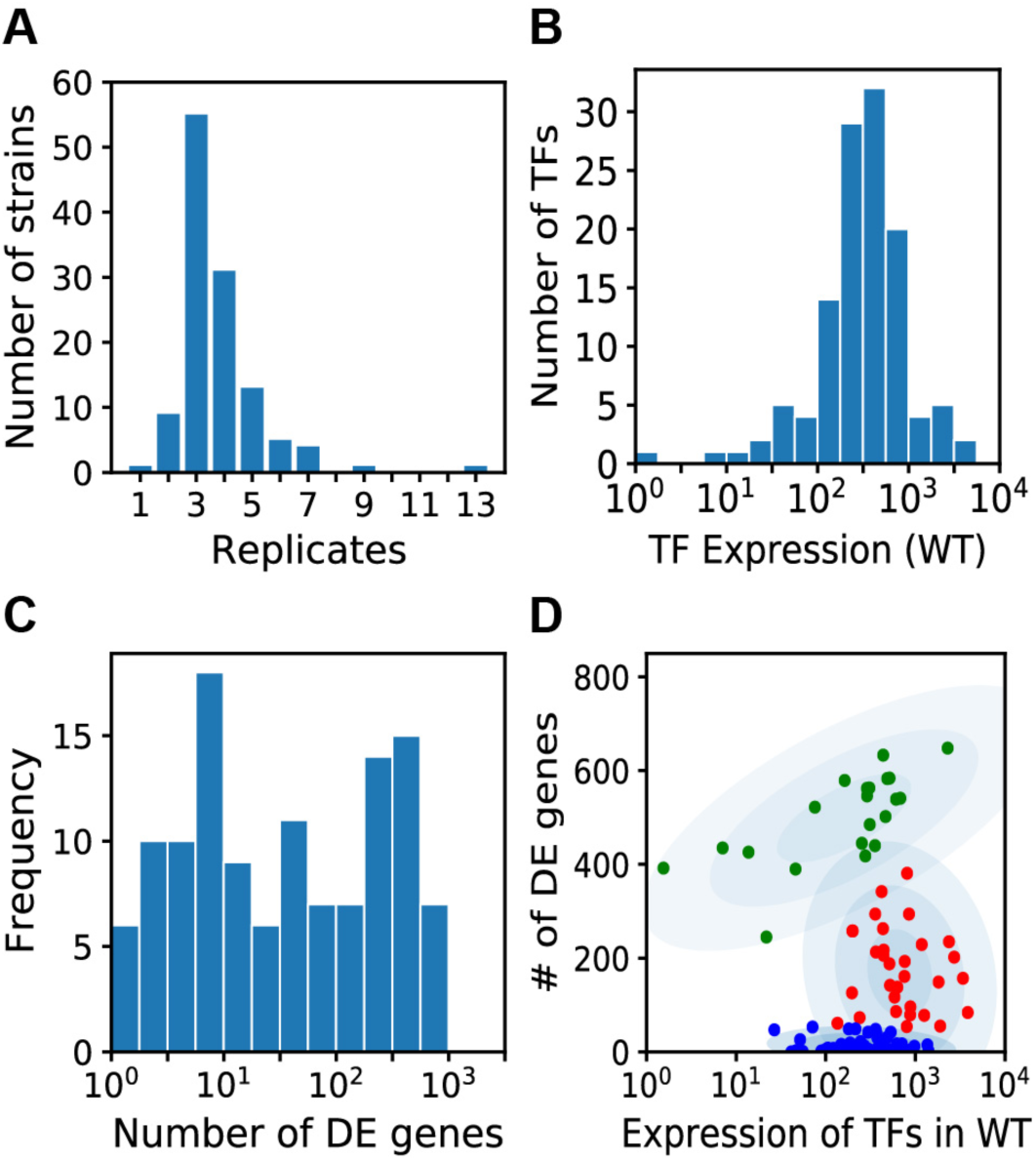
(A) Histogram of the number of RNA-seq biological replicates performed for TF deletion strains. (B) Histogram of the expression levels of TFs (normalized read counts) in WT cells. Only the 120 TFs that were deleted were included. (C) Histogram of the number of DE genes in TF deletion strains. (D) The number of genes differentially expressed in response to TF deletions as a function of WT expression levels of the deleted TFs. The red cluster was enriched for TFs implicated in capsule formation.

Next, for each TF deletion strain we calculated the number of differentially expressed (DE) genes, which may be functional targets of the TF (Fig. 1C and Supplementary Fig. S1B). A gene was defined as DE if the adjusted P-value of expression in the deletion versus WT was below 0.01 and the absolute log_2_ fold-change (FC) relative to WT was greater than 0.4 (https://doi.org/10.5281/zenodo.17193388 for non-TF-encoding genes and supplementary File S3 for TF-encoding genes). Half of the TF-deletion strains had 1-20 DE genes, suggesting that they have relatively specific functions in our conditions, while the other half had more than 20. Overall, the expression level of a TF in the WT strain was not correlated with the number of genes that were DE in the TF deletion (Pearson correlation *ρ* = 0). However, when we plotted the number of genes that responded to a TF deletion against that TF’s WT expression level, TFs clustered into three groups (Fig. 1D). TFs in the blue cluster regulated few to no genes, despite the broad range of expression levels of those TFs in WT cells. In the green cluster, the expression level of each TF correlated with the number of genes that were DE when that TF was deleted (Pearson correlation *ρ* = 0.57, *P* < 0.01). TFs in the red cluster were expressed at a relatively high level and had an intermediate number of DE genes (54-381). This group was enriched for TFs where the corresponding deletion strain has an abnormal capsule phenotype (Hypergeometric p = 0.0007), whereas the other clusters were not (hypergeometric p = 0.97 and p = 0.98; see Supplemental Methods for details).

### A TF network map for *Cryptococcus neoformans*

We used NetProphet3 (Abid and Brent, 2023) to construct a TF network map for *C. neoformans*. The goal of NetProphet3 is to predict direct and functional TF-target relationships, in other words, those in which a TF binds the promoter of a target gene and thereby influences its expression. It achieves this by using a machine learning (ML) algorithm to learn patterns in gene expression data that are indicative of direct regulation. ML algorithms are first trained to make predictions on cases where the true answer is known. They are then applied to make predictions where the true answer may not be known. During training, NetProphet3 uses available TF binding location data, such as from ChIP-Seq experiments, to indicate whether binding of the TF to a gene promoter is known to occur; this is assigned as a label, with a value of 1 (binding) or 0 (no binding) (Fig. 2A). In the prediction phase this information is no longer available, so only the expression levels of a gene after a TF perturbation are used to predict whether the TF binds its promoter region. In both phases, the gene expression data used to make predictions is first processed in various ways, producing the actual inputs to NetProphet3 (called features). Specifically, the features include the fold-change in expression of the target when the TF is perturbed (DE feature) and the relationships between the expression of TFs and the expression of target genes across all gene expression samples (LASSO and BART features; see Methods).

**Fig 2.**
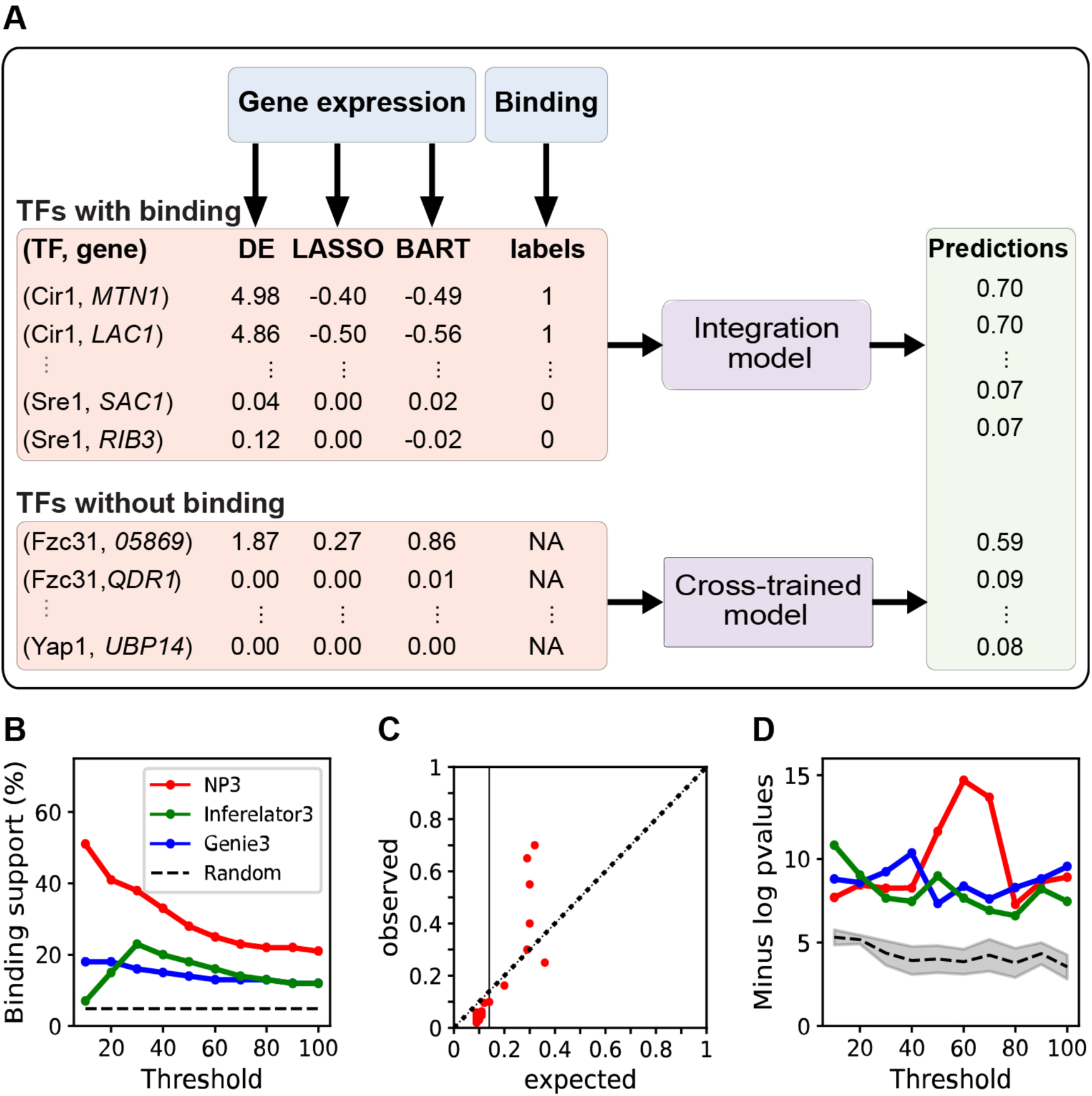
(A) Diagram of the NetProphet3 data flow, as described in the text. (B) Evaluation of the cross-trained mode, where Cryptococcus binding location data is not used in training, for agreement with binding data. Only TF-target edges where binding data is available for the TF were evaluated. Shown is the performance of NetProphet3 (red) versus random expectation (black dashed line) and two other established methods for inferring TF networks from gene expression data. Threshold, the number of highest-scoring edges included in the network, divided by the number of TFs. (C) ChIP-Seq validation of NetProphet predictions. Expected, NetProphet score (predicted probability); observed, actual validation of predictions by ChIP-seq data. (D) Evaluation of the NetProphet3 network after integrating binding data, using the GO metric; y-axis values plotted are the median across all TFs of the -log p value of the most significant GO term for that TF. The other algorithms indicated in panel (B) are shown for comparison. Shaded area, 95% confidence interval.

When the trained NetProphet3 model is applied to gene expression data, it produces a binding probability for each TF-gene pair (Fig. 2A, Predictions), which is interpreted as the probability that the gene is a direct, functional target of that TF. The predicted probability is high if NetProphet3 identifies patterns in the gene expression data that suggest the TF binds in the gene’s promoter. Since very little ChIP-seq binding location data is available for *Cryptococcus*, we trained NetProphet3 on *S. cerevisiae* binding and expression data (see Methods for details). We refer to this as NetProphet3’s *cross trained* mode. We then used this trained model on our gene expression data from *C. neoformans* TF deletion strains (without binding location data). The model is trained to combine evidence from features to make reasonable probability estimates. Importantly, the features do not include the identity of specific TFs or target genes. We therefore hypothesized that it would perform as well on *C. neoformans* data as it did on *S. cerevisiae* data.

### The *C. neoformans* network map consists of direct interactions

We next used two evaluation metrics to assess whether high scoring TF-target edges in our map are direct and functional. One of these is based on binding data and one is based on gene function annotation. For the first one, we defined a binding evaluation metric as the fraction of predicted TF-target edges that are supported by physical binding data (Abid and Brent, 2023). We used this to evaluate the network, which had been trained using only *S. cerevisiae* data and then applied to our *Cryptococcus* gene expression data. We performed this evaluation for 10 *C. neoformans* TFs with associated physical binding data: Cir1 (Do, et al., 2020), Cuf1 (Garcia-Santamarina, et al., 2018), Gat201 (Homer, et al., 2016), Gat204 (Homer, et al., 2016), Hap-X (Do, et al., 2020), Nrg1 (Gish, et al., 2016; Maier, et al., 2015), Liv3 (Homer, et al., 2016), Pdr802 (Reuwsaat, et al., 2021), Sre1 (PRJNA557210), and Usv101 (Gish, et al., 2016; Maier, et al., 2015).

True TF-target edges represent a small fraction of the universe of possible TF-gene edges. For this reason, we evaluated only the top scoring TF-gene edges. To do this we first ranked all TF-gene edges where the TF was one of the 10 that had binding data, from highest to lowest scores. We then generated networks that included a total number of edges equal to various multiples of the number of TFs. The most stringently thresholded network retained only 100 edges, equal to ten times the number of TFs with binding data (on average 10 per TF, although different TFs have different numbers of targets). Finally, we plotted the fraction of edges with binding support for networks of various sizes (Figure 2B).

Because we ranked edges by score, the networks with fewer targets per TF had higher average edge scores and were expected to have greater binding support. Indeed, for the smallest network, over 50% of predicted edges were supported by binding data, compared to a random expectation of 4% (Fig. 2B). Since ChIP-seq studies can miss true interactions, this is an excellent outcome and gives us high confidence in the remaining predictions.

As expected, increasing the network size led to a reduction in the fraction of edges supported by binding data (Fig. 2B). This shows that higher scoring edges are more likely to be direct. Furthermore, the performance of the *C. neoformans* network with the binding metric is comparable to that of the *S. cerevisiae* network produced by NetProphet3 (Abid and Brent, 2023), indicating that the NetProphet3 method developed using *S. cerevisiae* is equally applicable to *C. neoformans*. Since no *Cryptococcus* binding data were used at any point in the construction of the network, these are effectively prospective experiments, providing a good estimate of accuracy.

For comparison, we also ran two popular network inference algorithms from other labs, Genie3 (Huynh-Thu and Geurts, 2018) and Inferelator3 (Skok Gibbs, et al.) on our data (Fig. 2B). The results showed that NP3 substantially outperformed both of these in terms of ChIP-Seq support.

The TF-target edge scores output by NP3 are intended to be estimates of the probability that the target’s promoter would be bound by the TF in a ChIP-seq experiment. To test whether these scores were reasonable estimates, we selected sets of edges within various probability score ranges and calculated both the expected validation rate for each set (the average of its edge scores) and its actual validation rate, based on ChIP-Seq data (Fig. 2C). The vertical line shows the minimum score for inclusion in the network of 100 targets per TF. Above this score threshold, the observed rate was either very close to the expected rate or above it, indicating that the scores are reasonable estimates of the validation rate and in some cases are even too pessimistic.

### Incorporating Cryptococcus binding data into the network

Once evaluation with Cryptococcus binding data was complete, we produced a second network that incorporated the binding data into the network by using NP3’s integration mode (Abid and Brent, 2023) (see Supplemental Methods). The two networks differ only in the edges emanating from the 10 TFs for which we had ChIP-Seq data. The second network is used for the remainder of this paper and is also available at https://doi.org/10.5281/zenodo.17193243. This network map can be searched and visualized at http://cryptococcus.net.

### The *C. neoformans* network map is functionally coherent

The second evaluation metric we applied was based on gene ontology (GO) biological process annotations. This GO metric reflects the degree to which each TF’s targets are enriched for genes that share a common functional annotation and are therefore biologically coherent. For this evaluation, we downloaded *C. neoformans* annotations from the UniProt website, most of which were transferred from *S. cerevisiae* based on sequence similarity (see Methods for details). Overall, 799 GO biological processes were transferred to *C. neoformans*, annotating over 3,000 genes, and the target sets of 30 TFs in our network were significantly enriched for at least one GO biological process term (Supplementary Table S1). For each TF and each biological process annotation, we then calculated a hypergeometric p-value (see Methods for details). A significant p-value indicates that more of the TF’s target genes are annotated with the function than would be expected by chance, and hence the TF’s targets are functionally coherent. To evaluate statistical significance, we created 30 random networks by permuting TFs and targets and evaluated them using the same metric. Although there is some fluctuation in absolute level, the performance of our network significantly exceeded those of the random networks at every threshold, indicating that each TF’s targets in our network were much more functionally coherent than would be expected by chance (Fig. 2D). This further supports the validity of the network. The Genie3 and Inferelator3 networks had similar functional coherence, although NetProphet3 outperformed them on networks of intermediate size.

### The most influential TFs are enriched for those that are known to regulate capsule

For this study, we generated or obtained deletion strains lacking 120 of the 165 TFs in the Cryptococcus genome. We were unable to delete the other 45. However, NetProphet3 still predicts targets for these TFs. It does this by using non-DE features obtained by analyzing all gene expression profiles together (Fig. 2A, LASSO and BART; see Methods). We therefore had predictions for all 163 TFs that were expressed in our dataset, so our network included the 16,300 highest-scoring TF-target edges. 161 of the TFs had at least one target gene under the conditions assayed. The total number of regulated genes (4,103) corresponds to 60% of those that were measurably expressed in our studies and 49% of all *C. neoformans* genes. This network is the basis for our analyses below.

NetProphet3 outputs probability scores for all possible TF-gene edges, most of which have very low scores. When selecting a set of predicted edges for further analysis, there is a tradeoff between including only edges predicted with high confidence and including enough edges to characterize the organism’s network comprehensively. As a reasonable compromise, for further analysis we chose a set that included on average 100 targets per TF. This set consists of all edges scoring above 0.14 (Fig. 3A, dashed line).

**Fig 3.**
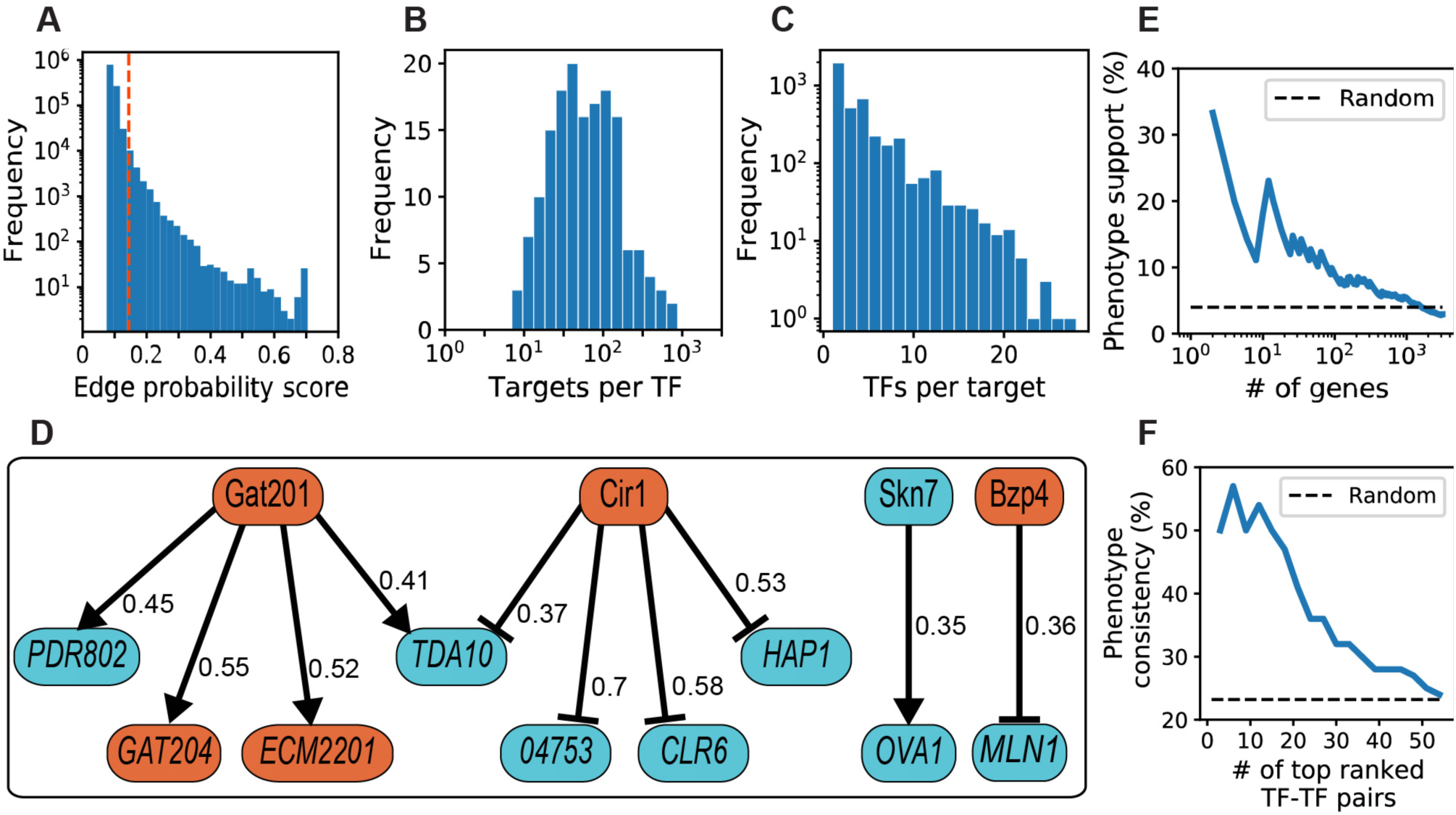
(A) NetProphet3 probability scores for all TF-gene edges. The vertical dashed orange line indicates the score threshold corresponding to 100 TF-target edges per TF on average. (B) The number of targets per TF. (C) The number of direct TF regulators per gene. (D) The top 10 high scoring TF-target edges in the capsule subnetwork. Colors indicate the phenotype of strains deleted for the corresponding genes: orange for hypocapsular and cyan for hypercapsular. Arrows denote activation and T-heads denote repression. All target genes except for *TDA10* and *CNAG_04753* encode DNA-binding proteins. The Gat201, Gat204 edge has been previously detailed (Chun, et al., 2011). (E) Genes were ranked by the number of capsule TFs that regulate them and the percentage of genes where the deletion strain exhibits a capsule phenotype was plotted for various cutoffs. (F) Percent of TF-TF pairs ranked above the indicated threshold that share a target and show phenotype consistency. Dashed line, percent of all TF-TF pairs that show phenotype consistency.

The more targets a TF has, the more influence it can potentially exert on the transcriptional state of the cell. In our network, the number of direct targets of each TF ranged from 7 to 884 (Fig. 3B). When we ranked TFs by the number of targets, the top 16 TFs (10%; Supplementary File S5) covered 6,709 (41%) of the 16,300 TF-target edges in our network. Surprisingly, these 16 TFs regulate 81% of the 4,103 genes in the network, showing their disproportionate impact. We were interested to notice that 10 of these TFs are known to regulate the capsule, based on the phenotypes of corresponding deletion strains (File S2). This is significantly enriched over the fraction of all TFs that are known to regulate the capsule (63% versus 31%; hypergeometric p = 0.006; see Discussion).

In addition to targets per TF, we examined the distribution of TFs per target gene (Fig. 3C). The distribution peaked at one regulator per target gene and dropped off exponentially, with one outlier gene having 55 regulators (CNAG_04126, not plotted in Fig. 3C). When we ranked genes by this metric, we found that the top 10% of targets (410) participated in 33% (5,315) of the TF-target edges. On average these top 10% targets have 13 regulators (Supplementary File S5). Also, these 410 targets were generally enriched for capsule-implicated genes (those whose deletion perturbs capsule; hypergeometric p < 3 × 10^−6^; Supplementary File S5).

Many TFs are both powerful regulators and highly regulated, thus occupying central positions in transcriptional networks. We defined hub TFs as those that are in the top 10% both by the number of TFs that regulate them and by the number of genes they regulate. Our network contained 14 hub TFs (Table 1), of which eight are capsule implicated (hypergeometric p = 0.007; see Discussion).

**Table 1.**
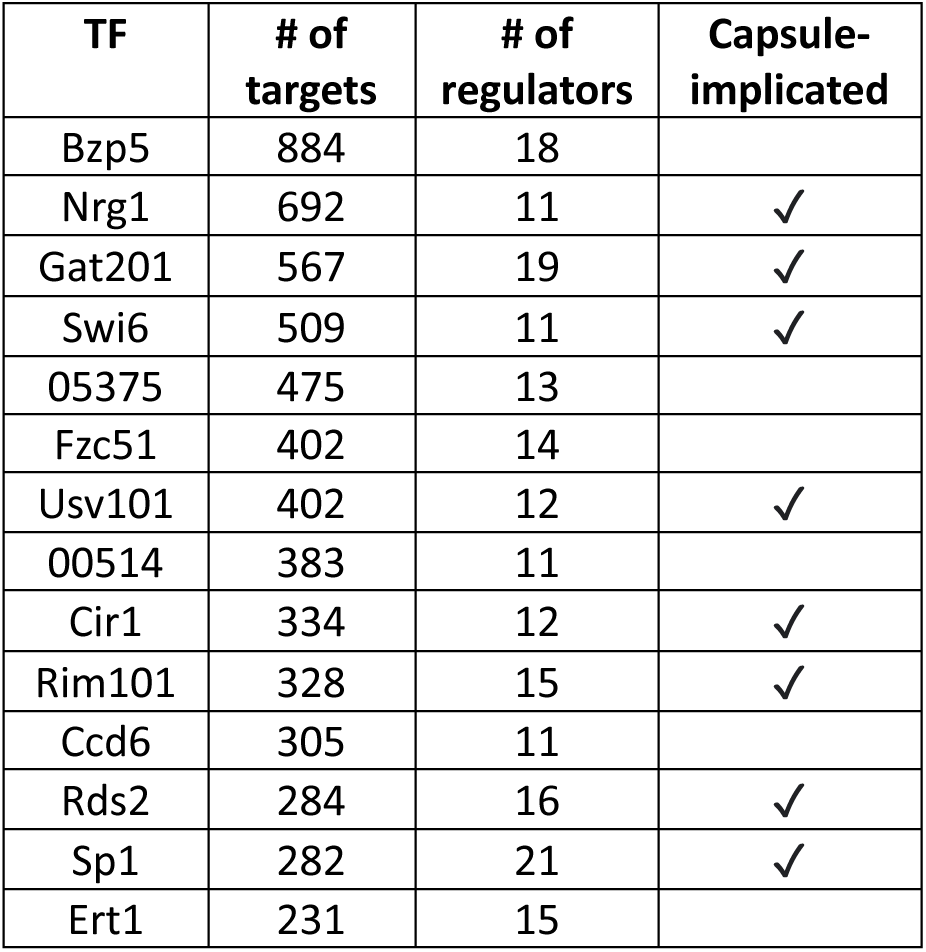
Hub TFs. We define hub TFs as those that are in the top 10% of TFs ranked by number of direct targets and are encoded by a gene in the top 10% of genes ranked by number of regulators. CNAG_05375 is an unnamed gene.

| TF | # of targets | # of regulators | Capsule-implicated |
| --- | --- | --- | --- |
| Bzp5 | 884 | 18 |  |
| Nrg1 | 692 | 11 | ✓ |
| Gat201 | 567 | 19 | ✓ |
| Swi6 | 509 | 11 | ✓ |
| 05375 | 475 | 13 |  |
| Fzc51 | 402 | 14 |  |
| Usv101 | 402 | 12 | ✓ |
| 00514 | 383 | 11 |  |
| Cir1 | 334 | 12 | ✓ |
| Rim101 | 328 | 15 | ✓ |
| Ccd6 | 305 | 11 |  |
| Rds2 | 284 | 16 | ✓ |
| Sp1 | 282 | 21 | ✓ |
| Ert1 | 231 | 15 |  |

### The *C. neoformans* network map is biologically coherent

We took advantage of the abundant available data on capsule phenotypes to assess whether the *C. neoformans* network makes biological sense. For this, we first created a capsule subnetwork composed of all TF-target edges for which both the TF and target are known to influence capsule phenotype. This capsule subnetwork includes 113 target genes, of which 49 encode TFs, linked by 318 TF-target edges.

Focusing on the capsule subnetwork, we investigated whether the predicted direction of regulation (activation or repression) at each edge was consistent with the capsule phenotypes of a TF and its target gene. We considered an edge consistent when (1) the TF activates the target, and both demonstrate the same capsule phenotype upon deletion (hypercapsular or hypocapsular) or (2) the TF represses the target and they have opposite capsule phenotypes. In Figure 3D, we show the 10 highest-scoring TF-target edges of the subnetwork; notably, many of the targets themselves encode TFs. All the edges shown were consistent with the capsule phenotype (binomial p = 0.02) except for two: Gat201-*PDR802* and Gat201-*TDA10*. In the case of *TDA10*, the negative regulation by Cir1 may be stronger than the positive regulation by Gat201; it cannot conform with both.

We speculated that target genes regulated by the highest number of capsule-implicated TFs were most likely to have a capsule phenotype upon deletion. To test this, we ranked genes by the number of capsule TFs that regulate them from highest to lowest and calculated, at different thresholds, the percentage of genes that have a capsule phenotype (Fig. 3E). We found that genes regulated by the highest number of capsule TFs were much more likely to be capsule genes than those that were regulated by a lower number of capsule TFs: 9 out of the top 100 genes in this ranking were capsule-implicated, while only 2% of all genes are capsule-implicated (hypergeometric p = 0.02).

### TF-TF pairs that share many targets

We previously showed that TFs that share large fractions of their targets are more likely to work together as a physical protein complex than those that do not (Abid and Brent, 2023). TF-TF pairs that share many targets but do not form a physical complex may cooperate by other mechanisms, such as being co-regulated by a third factor or binding the same DNA sequences. To gain insight into the TF-TF relationships of capsule TFs, we calculated the Jaccard similarity between target sets of all capsule-implicated TF pairs in the network (Supplementary File S6). The Jaccard similarity is the number of targets in common divided by the total number of individual targets regulated by the two TFs. It is equal to one if all targets are shared by both TFs, and zero if none are shared. For each TF-TF pair, we determined whether this similarity was significantly higher than would be expected by chance (hypergeometric adjusted p < 0.05), indicating that the TFs are likely to cooperate (as defined above). For this analysis, we used only TF-target edges that had high NetProphet3 probability scores (greater or equal to 0.2). These TF-TF interactions form a second subnetwork, the capsule TF subnetwork, that includes 43 capsule TFs linked by 56 TF-TF edges.

We observed that these TF pairs tended to regulate their targets in the same direction if the two TFs had the same capsule phenotype and in the opposite direction if the TFs had opposite capsule phenotypes. We considered the TF-TF pair to display phenotype consistency if both TFs had the same capsule phenotype and at least 80% of their shared targets were regulated in the same direction, or if they had opposite phenotypes and at least 80% of their shared targets were regulated in the opposite direction. We ranked these TF-TF pairs by the significance of their Jaccard similarity and calculated the fraction of pairs above different rank thresholds that display phenotype consistency (Fig. 3F). This fraction was much higher than would be expected at random and tended to decline as the significance of their target-set overlap declined, consistent with the notion that pairs with the highest target overlap are most likely to work in a coordinated fashion.

Relationships between cryptococcal TFs implicated in capsule have been reported in several contexts, including some efforts to map regulatory interactions (Chun, et al., 2011; Gish, et al., 2016; Haynes, et al., 2011; Kumar, et al., 2011; Maier, et al., 2015). However, these have generally been limited to relatively small groups of genes and focus on direct regulation of one TF by another. Here we focus on the overlap of targets among all capsule TFs. Figure 4 shows all pairs of capsule TFs that share many more targets in common than would be expected by chance (*p* < 10^−5^), along with the targets that they share. Notably, half of these TFs are hub TFs, highlighting their central role in transcriptional regulation. While most of the regulatory relationships are activating, several TFs (Usv101 and Cir1) are uniformly repressive in this context.

**Fig 4.**
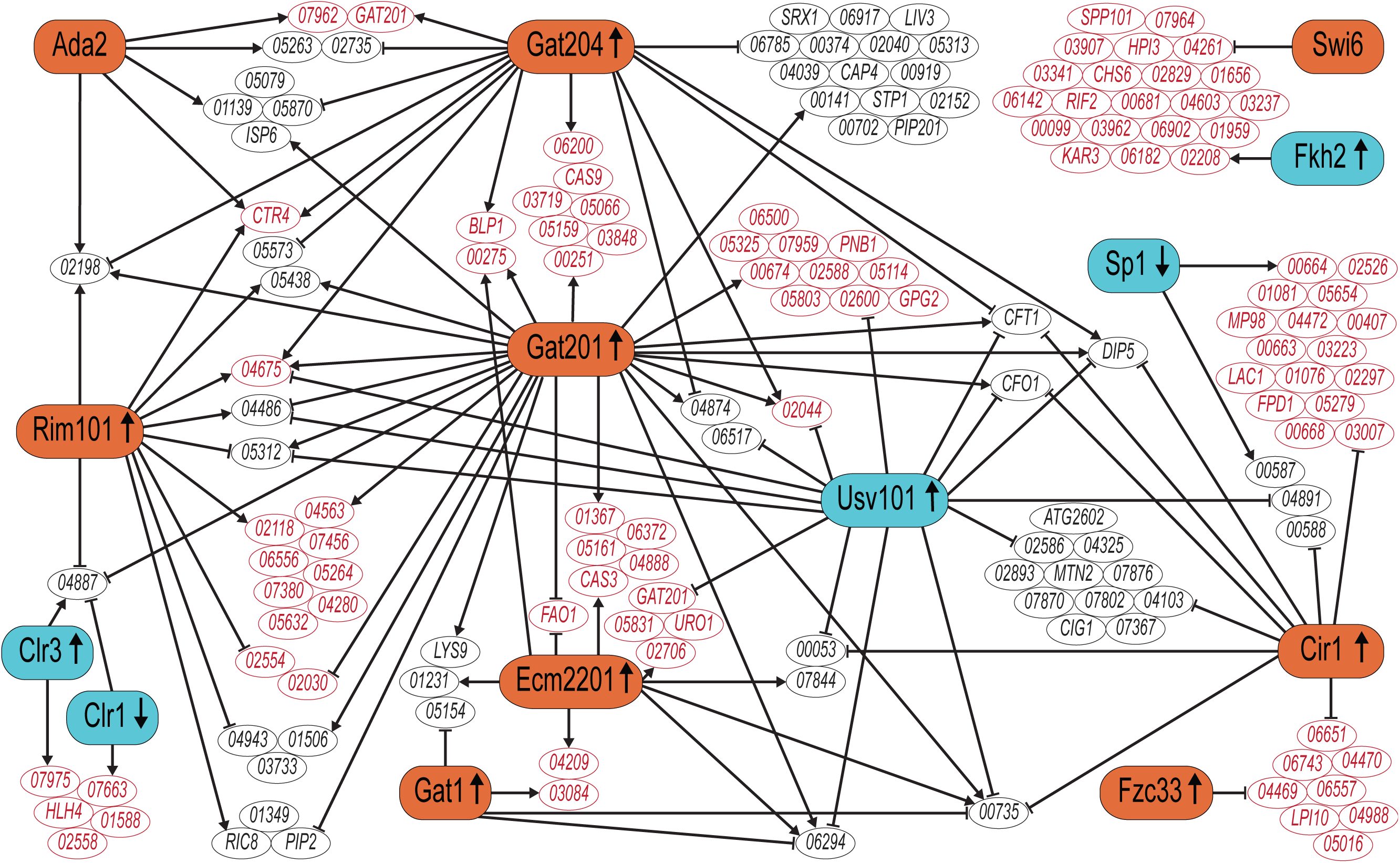
A TF network map of only shared targets of the 12 TF-TF pairs with the most significant hypergeometric p-values for their shared targets. These pairs are Gat201-Gat204, Usv101-Gat201, Swi6-Fkh2, Usv101-Cir1, Gat201-Rim101, Sp1-Cir1, Ecm2201-Gat201, Clr1-Clr3, Gat204-Ada2, Ecm2201-Gat1, Usv101-Ecm2201, Cir1-Fzc33. Orange and blue shading indicates that cells lacking the TF are hypocapsular or hypercapsular, respectively. TFs that are up-regulated and down-regulated in capsule inducing conditions are indicated by upward and downward arrows, respectively. Targets whose regulators display phenotype consistency (defined in main text) are labeled in red.

### Comparison of the *Cryptococcus neoformans* and *Saccharomyces cerevisiae* regulatory networks

Much of what we know about the functions and interactions of proteins in *C. neoformans* comes from studies of their orthologs in the model yeast *S. cerevisiae*. However, reconstruction of phylogenetic gene trees by sequence similarity comes with substantial uncertainty, and therefore identification of truly orthologous genes between species does, too. This is especially problematic for genes belonging to families that have substantial internal sequence similarity within each species, as is the case for many TFs. Even when two TFs are true historical orthologs, their functions may have diverged during evolution, so non-orthologous TFs may have more functional similarity. To identify TF pairs that were both historical orthologs and retained shared function, we combined sequence similarity with functional similarity, defined as regulating putatively orthologous target genes. We denote these pairs as (TF*_Cn_*, TF*_Sc_*).

Currently, the best available resource on homology between *S. cerevisiae* and *C. neoformans* proteins was compiled by Kelliher et al, who published a list of 4,572 homologous protein pairs based primarily on amino acid sequence similarity (Kelliher, et al., 2016). In this list, a few *C. neoformans* genes were mapped to more than one *S. cerevisiae* gene because *S. cerevisiae* genes include paralogs derived from a whole genome duplication event. We were curious as to whether the 63 pairs of TFs in this list, assigned by sequence homology, also showed functional similarity based on overlap of their target genes. For this purpose, we returned to the full NetProphet3 network for *S. cerevisiae* ((Abid and Brent, 2023); https://doi.org/10.5281/zenodo.17196637) and *C. neoformans*.

First, we calculated a Jaccard similarity for each (TF*_Cn_*, TF*_Sc_*) pair based on their homologous target genes. As above, higher Jaccard similarity for a (TF*_Cn_*, TF*_Sc_*) pair means that a higher fraction of the TFs’ targets encode homologous proteins. To determine whether the Jaccard similarity was significantly greater than what would be expected by chance, we calculated a permutation-based empirical P-value (see Methods). Surprisingly, only three (TF*_Cn_*, TF*_Sc_*) pairs from the Kelliher et al sequence-based list had significantly similar target gene sets: (Mig1*_Cn_*, Mig1*_Sc_*), (Skn7*_Cn_*, Skn7*_Sc_*), and (Gat1*_Cn_*, Gat1*_Sc_*). None of the other pairs on this list had significant target-set overlap, despite solid sequence similarity (Supplementary File S7-A). Next, we examined the seven homologous pairs from the Kelliher list where both the *C. neoformans* and *S. cerevisiae* TF targets were enriched for a GO biological process (Table 2). Only one pair, (Fkh2*_Cn_*, Fkh2*_Sc_*), had overlap of their most significant GO terms, suggesting that the remaining pairs primarily regulate distinct biological processes.

**Table 2.** (TF*_cn_*, TF*_sc_*) pairs proposed by Kelliher et al. where the targets of each TF are enriched for a GO biological process term. Identity, amino acid identity from BLASTP; GO_cn_ and GO_sc_, most significant GO terms for the target sets in *C. neoformans* and *S. cerevisiae* networks, respectively.

| TF<sub>Cn</sub> | TF<sub>Sc</sub> | Identity | *Cryptococcus* GO term | *Cerevisiae* GO term |
| --- | --- | --- | --- | --- |
| Fkh2 | Fkh2 | 47% | cell cycle and DNA replication | regulation of DNA replication |
| Fzc1<br>2 | Mal33 | 40% | regulation of cell cycle and<br>DNA replication | carbohydrate transport |
| Mln1 | Rtg3 | 30% | carbohydrate transport | TCA cycle, amino acid and<br>nitrogen metabolism |
| Rsc8 | Swi3 | 36% | transcriptional regulation | transport of ubiquitinated<br>proteins |
| Swi6 | Swi6 | 25% | regulation of cell cycle | cell wall and meiosis |
| Yap1 | Yap1 | 48% | transcriptional regulation | response to oxidative stress,<br>sterol biosynthesis, response to<br>metal ion, sulfur metabolism |
| Zfc2 | Usv1 | 46% | peptidyl-lysine modification<br>to peptidyl-hypusine | metabolism of glycogen and<br>other carbohydrates |

In the previous analyses, we began with TF sequence similarity and looked for evidence of functional similarity. Next, we began with TF pairs that had similar target sets and asked whether the TFs had similar protein sequences. For this we identified (TF*_Cn_*, TF*_Sc_*) pairs based solely on overlap of their targets and selected those with the highest Jaccard similarity Index (>0.1). Surprisingly, BLASTP did not identify sequence similarity between most of these TFs, with only one pair (Gat201*_Cn_*, Gat4*_Sc_*), showing a significant alignment (evalue<1.0; Supplementary File S7-B). In general, pairs that shared homologous targets, and hence are more likely have similar functions, did not have similar protein sequences.

We next looked for (TF*_Cn_*, TF*_Sc_*) pairs that had a balance of both sequence and functional similarity, using less stringent cutoffs. First, we ran BLASTP to align all possible (TF*_Cn_*, TF*_Sc_*) pairs. 161 Cryptococcus TFs were in at least one pair with amino acid identity greater than 30%, 162 were in at least one pair with BLASTP e-value less or equal than 1, and 24 were in a pair that had a significant Jaccard similarity index (corrected empirical P < 0.05; see Methods). Seventeen Cryptococcus TFs were in a pair that met all these criteria. For each of these, we calculated a score that combined their sequence and functional similarity (SF score, see Methods). For each *Cryptococcus* TF, we selected the (TF*_Cn_*, TF*_Sc_*) pair with the highest SF score. If more than one TF*_Cn_* mapped to the same TF*_Sc_*, the TF*_Cn_* with the higher SF score was paired with that TF*_Sc_*, while the TF*_Cn_* with the lower SF score was mapped to its TF*_Sc_* with the second-highest score.

The 17 TF pairs that we identified across these widely diverged species are derived from the same common ancestor (based on sequence similarity) and retain shared functions (based on target sets). Interestingly, the cryptococcal TFs in nine of these pairs (Table 3) do not appear in the sequence-based list of Kelliher et al. The remaining eight were included in that list, but our analysis suggests that they pair best to different partner TFs in *S. cerevisiae* with high sequence and functional similarity (Table 4). For only two TFs, Gat1 and Mig1, do both approaches agree (see Discussion).

**Table 3.** Nine (TF*_Cn_*, TF*_Sc_*) pairs identified by our method based on sequence and functional similarity, but not by Kelliher et al., who used sequence similarity alone. Identity and e-value from BLASTP; Jaccard similarity and empirical P-values calculated as described in Methods.

| TF<sub>Cn</sub> | TF<sub>Sc</sub> | Identity (%) | E-value | Jaccard similarity | Empirical P-value |
| --- | --- | --- | --- | --- | --- |
| Gat201 | Gat4 | 41 | 2.16E-05 | 0.11 | 0 |
| 01069 | Ppr1 | 50 | 6.36E-05 | 0.10 | 0.01 |
| Fzc45 | Pip2 | 34 | 0.007 | 0.09 | 0.01 |
| Stb4 | War1 | 32 | 0.026 | 0.09 | 0 |
| Cir1 | Gzf3 | 59 | 2.09E-18 | 0.08 | 0 |
| Fzc19 | Tbs1 | 30 | 0.83 | 0.08 | 0.03 |
| Sip401 | Dal81 | 32 | 0.76 | 0.08 | 0 |
| Fzc44 | Mal33 | 52 | 0.17 | 0.07 | 0.04 |
| 06097 | Pdr3 | 48 | 7.40E-02 | 0.07 | 0.02 |

**Table 4.** (TF*_Cn_*, TF*_Sc_*) pairs identified using both our method based on sequence and functional similarity and the Kelliher et al. method based only on sequence similarity alone. Identity, e-value, Jacquard similarity, and empirical p-value derived as in Table 3.

| TF<sub>Cn</sub> | TF<sub>Sc</sub> | Identity | e-value | Jaccard similarity | Empirical P-value | Our method | Kelliher et al. |
| --- | --- | --- | --- | --- | --- | --- | --- |
| Ert1 | OAF1 | 41% | 1.86E-06 | 0.07 | 0.00 | ✓ |  |
| Ert1 | Ert1 | 49% | 2.82E-09 | 0.01 | 1.00 |  | ✓ |
| Fzc33 | Rgt1 | 33% | 0.046 | 0.07 | 0.01 | ✓ |  |
| Fzc33 | Sef1 | 43% | 7.69E-07 | 0 | N/A |  | ✓ |
| Gat1 | Gat1 | 70% | 3.12E-22 | 0.09 | 0.00 | ✓ | ✓ |
| Mig1 | Mig1 | 68% | 8.48E-27 | 0.09 | 0.00 | ✓ | ✓ |
| Ppr1 | Mal13 | 47% | 0.061 | 0.07 | 0.05 | ✓ |  |
| Ppr1 | Ppr1 | 25% | 1.28E-07 | 0.04 | 0.61 |  | ✓ |
| Rim101 | ZAP1 | 31% | 5.93E-12 | 0.08 | 0.01 | ✓ |  |
| Rim101 | Rim101 | 40% | 7.69E-07 | 0.04 | 1.00 |  | ✓ |
| Skn7 | Skn7 | 51% | 5.18E-31 | 0.06 | 0.03 | ✓ | ✓ |
| Skn7 | Hms2 | 28% | 5.48E-08 | 0.03 | 1.00 |  | ✓ |
| Usv101 | Msn2 | 59% | 1.41E-11 | 0.1 | 0.00 | ✓ |  |
| Usv101 | Usv1 | 62% | 3.59E-20 | 0.07 | 0.09 |  | ✓ |
| Usv101 | Rgm1 | 58% | 2.15E-21 | 0.05 | 0.99 |  | ✓ |

Of the 165 Cryptococcus TFs, fewer than half (72) have been mapped to *S. cerevisiae* TFs based on either our method or sequence similarity alone. The remaining 93 have apparently diverged and may serve functions that do not exist in *S. cerevisiae*. Interestingly, 29 of these 93 highly diverged TFs have target sets enriched for genes that do not have homologs in *S. cerevisiae* (FDR = 0.05; Supplementary File S8), thus highlighting the unique biology of *C. neoformans* compared to model yeast.

## Methods

### Cell growth

To maximize reproducibility of RNA-seq, we followed strictly controlled protocols for cell recovery from frozen stocks, initial culture in YPD, inoculation into preconditioned media, and growth (see Supplementary Methods) and (Kang, et al., 2025).

### RNA-Seq

RNA isolation, library preparation, and sequencing were by standard methods as detailed in the Supplementary Methods and (Kang, et al., 2025).

### NetProphet3

We processed counts of measured gene expression levels into normalized log_2_ fold changes using DESeq2 (Love, et al., 2014). Samples for each TF deletion strain were compared to all 122 WT samples taken together. We then ran NetProphet3; see (Abid and Brent, 2023) for details of how NetProphet3 works. Our dataset included perturbation profiles for 120 TFs. For those TFs, the differential expression (DE) feature for each gene was the log_2_ fold change of the gene’s expression in the TF deletion mutant, relative to its expression in the wild-type strain. For Gat204 and Liv3 we used TF perturbation data from Homer et al (Homer, et al., 2016). For the remaining TFs, which did not have DE data, we assigned all potential target genes a value of 0 (https://doi.org/10.5281/zenodo.17193620).

LASSO (Efron, et al., 2004; Friedman, et al., 2008) and BART (Chipman, et al., 2010; He, et al., 2019) are regression algorithms which were used to derive additional features for each potential TF-target edge by training models to predict the expression of each gene from the expression levels of all TFs. The LASSO feature for a TF-gene edge is the linear regression coefficient learned for that TF in the model for that gene. The BART feature for a TF-gene edge is the predicted change in expression of the gene when the TF’s expression changes from its highest to lowest value found in the data (see (Abid and Brent, 2023) for details).

As described in the text, we first trained NetProphet3 on binding and expression data from *S. cerevisiae* (NetProphet3 cross-trained mode, described in (Abid and Brent, 2023)). This training is not TF-specific; NetProphet3 simply learns patterns of gene expression features that are characteristic edges supported by binding location data. For the ten TFs that have binding location data in Cryptococcus, we trained a separate, TF-specific model that integrates binding and expression data (NetProphet3 integration mode, described in (Abid and Brent, 2023)).Finally, we combined networks produced by the cross-trained mode and the integration mode into one final network (Figure 2A).

### Evaluation metrics

#### Binding metric

We used available binding data, which was for only 10 TFs (see Results and Table S2), which contained 3,308 supported TF-target edges. Since these datasets and the information provided with them were so heterogeneous, we used different P-values for thresholds for each dataset (Supplementary Table S2).

#### GO metric

We used the GO-Term-Finder package (Boyle, et al., 2004) to calculate whether the targets of each TF were enriched for any GO biological process terms (Abid and Brent, 2023). GO annotations were downloaded from the UniProt website (Supplementary File S4) and transferred from *S. cerevisiae* annotations based on sequence homology. We then calculated a Hypergeometric P-value for enrichment of each TF’s targets with each GO biological process term. For TFs with at least one significant GO term, we calculated the median, across all significant GO terms, of their minus log P-value.

### Empirical P-value for Jaccard similarity between Cryptococcus and Saccharomyces TFs

We constructed 1,000 randomized Cryptococcus networks by shuffling confidence scores of TF-target edges. For each randomized network, we calculated the Jaccard similarity index for each TF-TF pair. For each TF-TF pair in the original network, the fraction of TF-TF pairs in randomized networks with a greater or equal Jaccard similarity is the empirical P-value.

### Sequence-functional (SF) similarity score calculation

For Cryptococcus TFs that aligned to an *S. cerevisiae* TF with at least 30% identity and an E-value of 1.0 or less, we compared their target sets and calculated an empirical P-value for the null hypothesis that the overlap is no larger than expected by chance (see previous paragraph). We then combined the alignment E-value with the empirical P-value for target overlap to obtain a sequence-functional similarity (SF) score via the following equation:

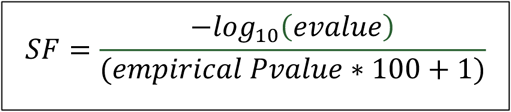

The SF is large when the alignment E-value is small (hence its negative logarithm is large) and the empirical P-value is small. The +1 in the denominator ensures that the fraction is still defined when the empirical P-value is zero.

## Discussion

A TF network map represents regulatory relationships by linking each TF to its direct, functional targets – the genes it regulates by binding to their regulatory DNA. This mechanistic approach stands in contrast to co-expression networks, which link pairs of genes that have correlated expression patterns. Co-expression networks have been constructed for Cryptococcus and used to identify new genes with a particular deletion phenotype, such as altered capsule thickness (O’Meara, et al., 2024). TF network maps have also been used to identify novel genes with a particular phenotype by using methods such as PhenoProphet (Maier, et al., 2015). In addition, TF network maps can be used for inferring changes in TF activity (Ma and Brent, 2021) and for determining which TFs regulate many of the same target genes, and hence may form a physical complex (Abid and Brent, 2023).

For *S. cerevisiae*, a wealth of comprehensive genomic resources has contributed to our understanding of gene regulation. There are multiple binding location datasets for most yeast TFs, obtained by using ChIP-chip (Harbison, et al., 2004), ChIP-exo (Rossi, et al., 2021), ChEC-seq (Lupo, et al., 2023; Mahendrawada, et al., 2023) and Transposon Calling Cards (Kang, et al., 2020). These are complemented by multiple comprehensive datasets, in which gene expression was profiled before and after each yeast TF was perturbed by gene deletion (Hu, et al., 2007; Kemmeren, et al., 2014), gene over-expression (Hackett, et al., 2020), or rapid protein degradation (Mahendrawada, et al., 2025). We previously used some of these data sets and our mapping algorithm NetProphet3 to build a comprehensive TF network map for *S. cerevisiae* (Abid and Brent, 2023). We wished to do the same for *C. neoformans*, but no comprehensive binding or TF perturbation datasets existed. To fill this gap, we created a comprehensive TF perturbation response dataset by carrying out RNA-Seq on strains in which the gene encoding a single TF had been deleted, a resource that we expect will be useful to the Cryptococcus research community. This dataset is completely independent of our large RNA-Seq dataset on responses to host-like conditions (Kang, et al., 2024).

With this dataset in hand, we used NetProphet3 to build a TF network map for Cryptococcus. To determine its reliability, we used two primary metrics. The first tested whether high-scoring edges (those NetProphet3 is most confident of) tend to be supported by ChIP-seq data where such data are available (Fig. 2B, 2C). By this metric, NP3 outperformed two other popular network inference algorithms that we applied to our gene expression data. The second metric tested whether TFs’ target sets are enriched for genes with shared biological functions at a rate higher than would be expected by chance (Fig. 2D). Both assessments indicated that our network is reliable.

Our previous *Cryptococcus* TF network map (Maier, et al., 2015) was built using gene expression data on only 41 TF-deletion strains. The current network uses data from 79 additional TF deletion strains, so it has more high-scoring edges from the new TFs. By our measures, the new network is significantly better (Fig. S2). We also built a website for the new network where users can visualize subnetworks comprising TFs and genes of interest (https://cryptococcus.net). We anticipate this will be of use to researchers in the field as they develop and test mechanistic hypotheses about cryptococcal TFs.

Capsule-implicated TFs played a central role in our network, comprising 8 of the 14 hub TFs (Table 1), significantly more than expected by chance. Surprisingly, no TF had capsule-implicated genes as more than 36% of its targets. Even for major capsule regulators such as Gat201, Cir1, and Nrg1, only 4-5% of their targets were capsule-implicated genes. This suggests the unexpected idea that there may not be any capsule-specific TFs. Instead, capsule genes appear to be regulated by TFs that also regulate many other processes. Large polysaccharide capsules are phylogenetically restricted among fungi, presumably evolved to manage species-specific challenges. Thus, one might expect that they could be deployed specifically in response to such stresses, independent of phylogenetically ubiquitous stress responses. However, our network suggests that this is not the case. Instead, it may be better to think of capsule as one of a large set of tools that are deployed together in response to certain stressful conditions. Nonetheless, genes that are coordinately regulated by multiple capsule-implicated TFs (Fig. 4) may be involved in capsule development or other key stress-response functions and hence of interest for follow-up studies.

Cryptococcus genes are often named after their putative orthologs in *S. cerevisiae* and implicitly assumed to have similar function. Putative orthologs are identified by protein sequence similarity (Kelliher, et al., 2016). However, TFs typically consist of a small DNA binding domain (DBD) and large intrinsically disordered regions that contain no identifiable functional domains. These disordered regions tend to have little sequence conservation across long evolutionary distances. Therefore, TF alignments between distantly related species are often possible only within the DBD, which comprises a small fraction of TF protein sequence. Furthermore, it is increasingly understood that, while DBDs may define a core DNA binding motif, binding locations are significantly impacted by the disordered domains (Brodsky, et al., 2020; Mindel, et al., 2024). These observations suggest that inferring functional homology between TFs from sequence alone may be difficult and error prone. To complement sequence-based approaches, we attempted to determine functional homology by comparing *C. neoformans* and *S. cerevisiae* TFs according to how many putatively orthologous target genes they shared. By combining sequence homology and functional homology, we identified 17 pairs consisting of a Cryptococcus TF and an *S. cerevisiae* TF that we consider true functional orthologs. Cryptococcus TFs that are not paired to *S. cerevisiae* orthologs may have evolved species-specific functions and therefore regulate many target genes that do not have *S. cerevisiae* orthologs. Alternatively, they may have retained their function while their target genes diverged from *S. cerevisiae* to the point where their orthologs cannot be identified by sequence comparison. Interestingly, sequence homology alone had suggested no orthologs for nine of the cryptococcal TFs in our 17 pairs in a previous study (Table 3), while for the other eight, our method suggested different orthologs from those suggested by sequence similarity alone. Overall, the sequence-based method tended to agree with our method only when protein identity was very high (∼70%). This suggests that TF network maps may, in general, be important for identifying functional orthologs between highly diverged species.

We were able to construct a high-quality TF network map using a large new dataset we generated of gene expression profiles from strains lacking TFs (perturbation response data). However, more data resources would make the network map even better. Although our genome sequence analysis identified 165 putative transcription factors in the genome of *Cryptococcus neoformans*, we were able to obtain or construct deletion mutants for only 120 of them. Some of the remaining 45 have been added to the Madhani deletion collection since this work was carried out. The remainder may be essential (Billmyre, et al., 2024). For those that are essential, newly implemented methods in the field, such as inducible protein degradation, (Huang, et al., 2025), may be useful. Another, complementary resource that is needed to fully illuminate the regulatory network is a comprehensive TF binding location dataset, potentially generated using ChIP-exo, ChEC-seq, or Transposon Calling Cards (Mayhew, 2016; Yen, 2023). Future integration of such data into our TF network using NetProphet3 will yield an enhanced TF network map that will further advance research into this fascinating and important pathogen.

## Acknowledgments

We are grateful to Cynthia Ma and Yu Sung Kang for their willingness to answer questions about their previous analyses of Cryptococcus RNA-Seq data.

## Data availability

All RNA-Seq data on Cryptococcus TF deletion strains used in this paper are available through the NCBI Gene Expression Omnibus under accession GSE297962.

