## Supplementary material for "Mapping the transcriptional regulatory network of a fungal pathogen by exploiting transcription factor perturbation": Online Supplement

Supplementary files

**File_S1_TFs.xlsx**

A list of the 120 TFs that were deleted, the number of RNA-Seq replicates we have for each, and the 163 TFs that were used in the construction of the TF network map.

**File_S2_capsule_genes.tsv**

A list of 164 genes from FungiDB whose deletions have abnormal capsule phenotypes.

**File_S3_expr_reg_log2_fc_163_120.xlsx**

Differential expression of TF-encoding genes (Log_2_ fold change of TF deletion over wild-type). Each column provides data for a different TF deletion strain.

**File_S4_gene_go_association.txt**

GO annotations downloaded from the UniProt website and processed into a GAF file for input to the package GO-Term-Finder.

**File_S5_top_outdegree_top_indegree.xlsx**

Top 16 (10%) TFs that have the highest number of targets, and top 410 (10%) genes that have the highest number of regulators.

**File_S6_net_tf_tf.xlsx**

TF-TF pairs that have a Jaccard similarity greater than zero. The file includes the number of shared targets, Jaccard similarity index, p-value, and adjusted p-value.

**File_S7_tf_tf_pairs_mappings.xlsx**

A. (TF*_cn_*, TF*_sc_*) pairs that were mapped using the list of Kelliher et al. (Kelliher, et al., 2016)
B. (TF*_cn_*, TF*_sc_*) pairs that were mapped using homologous targets of *C. neoformans* and *S. cerevisiae*. If a Cryptococcus TF does not map to the S. cerevisiae TF at all, the columns “% identity”, “evalue”, and “qcov” are left blank.

qcov (query coverage) is the percentage of the query sequence that is covered by a BLASTP alignment with a sequence from the database.

**File_S8_tfs_not_mapped.xlsx**

Cryptococcus TFs that were not mapped to an *S. cerevisiae* TF by either our method or the Kelliher method. The file includes for each TF: the systematic name, the common name, whether it is known to regulate capsule, and the corrected p-value for enrichment with target genes that do not have *S. cerevisiae* homologs according to Kelliher et al. (Kelliher, et al., 2016).

Large files on Zenodo

The TF network map constructed using NetProphet3 cross-trained and integration modes.

<https://doi.org/10.5281/zenodo.17193243>

Differential expression of non-TF encoding genes (Log_2_ fold change of TF deletion over wild-type). Each column provides data for a different TF deletion strain.

<https://doi.org/10.5281/zenodo.17193388>

The DE network that was used to run NetProphet3. Rows are TFs and columns are target genes.

<https://doi.org/10.5281/zenodo.17193620>

The *S. cerevisiae* TF network map that was generated using NP3 10-fold cross validation (Abid and Brent, 2023) and used in this study. Format is tab separated values (.tsv).

<https://doi.org/10.5281/zenodo.17196637>

Supplementary Methods

Cell growth

Strains were streaked from -80 °C to YPD agar, grown at 30 °C for 2 days, and a single colony was inoculated into 4 ml yeast peptone dextrose (YPD) and grown overnight at 30 °C shaking at 230 rpm. One day before the experiment, 1 ml of this culture was inoculated into 100 mL YPD and grown as above. At the beginning of the experiment, this culture was spun down, washed in 25 ml room temperature (RT) PBS, and resuspended in 10 ml PBS. 2 x 10^8^ cells were then inoculated into a T75 flask (MIDSCI TP90076) containing 20 ml DMEM (Sigma D6429) and incubated at 37 °C under 5% CO_2_. Each experiment included WT and one or more TF deletion strains.

Flasks were harvested at 90 min after inoculation by transferring 18.5 ml of culture to a tube on ice containing 2 ml RNA Stop solution (5% tris-saturated phenol in ethanol). The cells were then sedimented (3000g, 5 min, 4 °C), the supernatant fraction decanted, and the pellet frozen in liquid nitrogen and stored at -80 °C.

RNA Isolation

Frozen cell pellets were thawed on ice and 875 ml TRIzol (Ambion 15-596-018) was added. Cells were then lysed by beating with 750 ml 0.5mm silica-zirconia beads for 3 min in a Minibeadbeater (Biospec Products) and RNA was isolated following the manufacturer’s protocol. Residual DNA was removed with the Turbo DNA-free Kit (Life Technologies AM1907) and polyA+ RNA was isolated from ~1 mg of total RNA using the NEBNext Poly(A) mRNA Magnetic Isolation Module.

Library preparation and sequencing

Libraries were constructed using the NEBNext Ultra Directional RNA Library Prep Kit from Illumina and samples were pooled at 10 nM. The pools were sequenced on a NextSeq 500 using the High75v2 kit as a 1x75 with 7 bp Index1 and 6 bp Index2 with 1% PhiX. Fastq files were demultiplexed using the Illumina bcl2fastq2 allowing 1 bp mismatch.

Enrichment for capsule phenotype

We first identified genes with reported capsule phenotypes from the FungiDB website ([www.fungidb.org](http://www.fungidb.org)). To assemble this list, we searched for the presence of the word ‘capsule’ in the ‘phenotype’ column of genes on this website and then excluded genes where the identified phenotype was described as normal (e.g. ‘normal capsule organization’, ‘normal width capsule’, and ‘normal capsule polysaccharide biosynthetic process’). The resulting list included 164 genes, of which 50 were TFs (Supplementary File. S2). Given any gene set, we determined whether it was enriched for capsule-implicated genes by using the hypergeometric test.

Data clustering for Fig. 1D.

For clustering, we used the Gaussian Mixture Model (GMM), a statistical model that represents data as a weighted combination of multiple Gaussian distributions, with each Gaussian representing a cluster. To fit this model, we employed the Expectation-Maximization (EM) algorithm, which requires specifying the number of clusters in advance. The software package used was: <https://scikit-learn.org/stable/modules/generated/sklearn.mixture.GaussianMixture.html#sklearn.mixture.GaussianMixture>

Integration mode

This mode is applied to predict scores for TF-target edges of the 10 TFs that have binding data available (Table S2). For each TF, NetProphet3 trains a separate XG-boost model using binding data for that TF and the gene expression data on all TFs. This TF-specific model, which is intentionally over-fit, integrates information from the binding data and the gene expression data. Once trained, the model is applied to the same gene expression data. Its predictions for each gene provide the NetProphet score for the edge to that gene from the TF the model was trained on.

LASSO and BART features

NetProphet3 employs two regression algorithms to generate evidence scores for TF-target edges: (a) Least Absolute Shrinkage and Selection Operator (LASSO) and (b) Bayesian Additive Regression Trees (BART). TF-target evidence scores are used as features in the NetProphet3 framework (Fig. 2A). The intuition behind using regression algorithms is that a correlation between mRNA levels of a gene and those of a TF suggest that the TF might regulate that gene. These algorithms work by regressing the mRNA levels of each target gene against those of all TFs. LASSO excels at linear feature selection, while BART captures non-linear interactions. Using both methods thus provides better results than using either one alone (Abid, 2023).

Supplementary tables

| **TF** | **GO-ID** | **Description** | **Corrected P-value** |
| --- | --- | --- | --- |
| Fkh2 | GO:0006260 | DNA replication | 1.72E-15 |
| Rim101 | GO:0016052 | carbohydrate catabolic process | 1.18E-03 |
| Sp1 | GO:0071852 | fungal-type cell wall organization or biogenesis | 1.58E-03 |
| Yrm103 | GO:0006812 | cation transport | 3.39E-03 |
| Stb4 | GO:0008643 | carbohydrate transport | 2.51E-04 |
| Swi6 | GO:0007049 | cell cycle | 4.72E-12 |
| Fzc25 | GO:0006183 | GTP biosynthetic process | 1.27E-04 |
| Ccd6 | GO:0042254 | ribosome biogenesis | 4.23E-30 |
| Fzc44 | GO:0008643 | carbohydrate transport | 2.25E-04 |
| Fzc19 | GO:0046349 | amino sugar biosynthetic process | 7.53E-04 |
| Yap1 | GO:0006357 | regulation of transcription by RNA polymerase II | 4.13E-04 |
| Fzc10 | GO:0006355 | regulation of transcription, DNA-templated | 5.25E-03 |
| Hob3 | GO:0006260 | DNA replication | 1.69E-10 |
| 03129 | GO:0042254 | ribosome biogenesis | 1.12E-35 |
| Fzc18 | GO:0006357 | regulation of transcription by RNA polymerase II | 3.80E-04 |
| Mal13 | GO:0045944 | positive regulation of transcription by RNA | 1.97E-03 |
| Fzc51 | GO:0042254 | ribosome biogenesis | 1.44E-08 |
| 05375 | GO:0042254 | ribosome biogenesis | 1.06E-19 |
| Mig1 | GO:0008643 | carbohydrate transport | 3.20E-03 |
| Mln1 | GO:0008643 | carbohydrate transport | 4.26E-05 |
| Hlh5 | GO:0006351 | transcription, DNA-templated | 1.08E-02 |
| Nrg1 | GO:0090329 | regulation of DNA-dependent DNA replication | 3.73E-07 |
| Fzc12 | GO:0030174 | regulation of DNA-dependent DNA replication | 1.09E-06 |
| Sip402 | GO:0034599 | cellular response to oxidative stress | 3.90E-03 |
| Gat5 | GO:0007049 | cell cycle | 1.35E-04 |
| Rsc8 | GO:0006355 | regulation of transcription, DNA-templated | 3.13E-03 |
| 04600 | GO:0045930 | negative regulation of mitotic cell cycle | 1.19E-02 |
| 05311 | GO:0006355 | regulation of transcription, DNA-templated | 1.49E-02 |
| Bzp4 | GO:0009226 | nucleotide-sugar biosynthetic process | 9.70E-03 |
| Zfc2 | GO:0008612 | peptidyl-lysine modification to peptidyl-hypusine | 1.09E-02 |

Table S1. TFs with target genes that were annotated by a GO biological term with a significant P-value. Gene IDs are available in File S1.

| **TF** | **# of bound genes** | **Paper/source** | **P-value**  **threshold** |
| --- | --- | --- | --- |
| Cir1 | 211 | (Do, et al., 2020) | Authors’ cut-off |
| Cuf1 | 398 | (Garcia-Santamarina, et al., 2018) | $<{10}^{-15}$ |
| Gat201 | 397 | (Homer, et al., 2016) | $<{10}^{-20}$ |
| Gat204 | 255 | (Homer, et al., 2016) | $\leq{10}^{-4}$ |
| Hap-X | 196 | (Do, et al., 2020) | Authors’ cut-off |
| Nrg1 | 505 | (Gish, et al., 2016; Maier, et al., 2015) | Authors’ cut-off |
| Liv3 | 245 | (Homer, et al., 2016) | $\leq{10}^{-3}$ |
| Pdr802 | 482 | (Reuwsaat, et al., 2021) | $\leq{10}^{-1100}$ |
| Sre1 | 109 | PRJNA557210 | Authors’ cut-off |
| Usv101 | 510 | (Gish, et al., 2016; Maier, et al., 2015) | Authors’ cut-off |

Table S2. The ten TFs that had binding (ChIP-Seq) data available at the time of writing the paper. In the last column, “authors’ cutoff” indicates that p values were not available, so all targets considered significant by the authors were included. In cases where p values were available, the largest threshold $\leq{10}^{-3}$so that no TF would have more than 500 bound targets was used and is indicated.

Supplementary figures

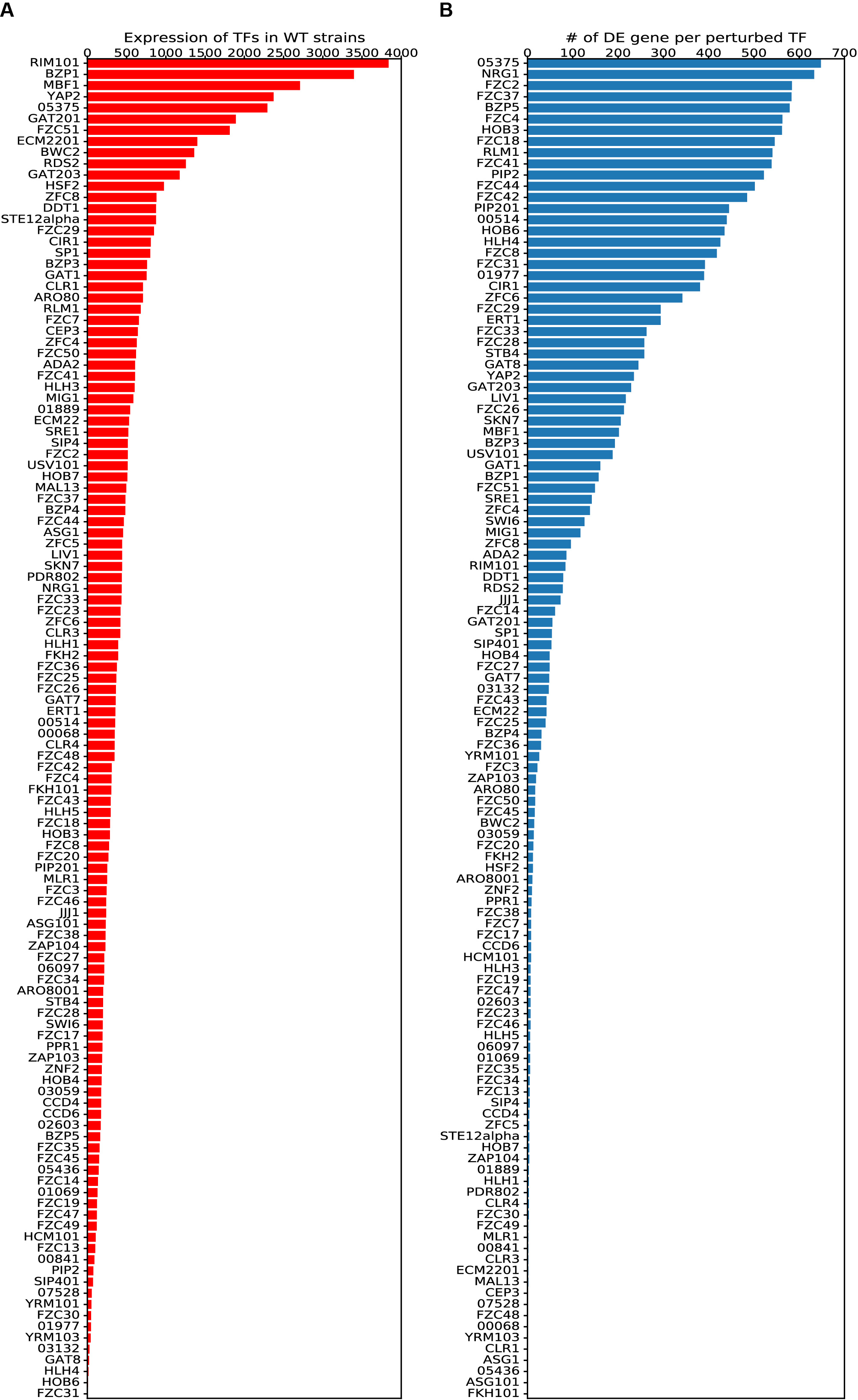

Fig. S1 (A). Normalized read counts for each deleted TF in a WT strain averaged across 122 biological replicates. (B) The number of DE genes after deleting each TF. The TF itself is not included in the DE genes.

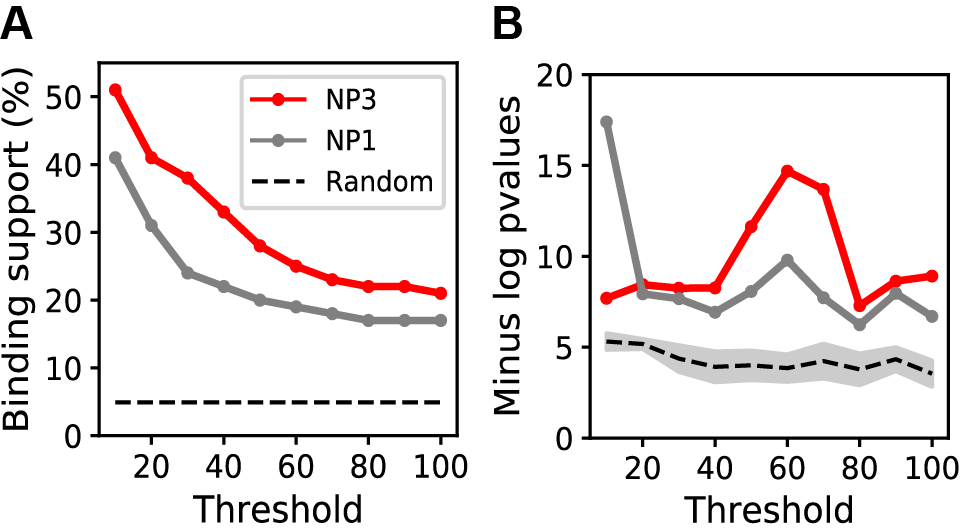

Fig. S2. Quality comparison of the NetProphet3 network built using the dataset reported here and the network described in (Maier, et al., 2015), which was built by applying the earlier NetProphet1 algorithm to gene expression data on only 41 TF deletion strains. (A) The new network (NP3) has more binding support at every score threshold. (B) The new network has greater functional coherence at every score threshold but one, a threshold which would yield a very small network.
